# Multiple inputs ensure yeast cell size homeostasis during cell cycle progression

**DOI:** 10.1101/226803

**Authors:** Cecilia Garmendia-Torres, Olivier Tassy, Audrey Matifas, Nacho Molina, Gilles Charvin

## Abstract

Coordination of cell growth and division is essential for proper cell function. In budding yeast, although some molecular mechanisms responsible for cell size control during G1 have been elucidated, the mechanism by which cell size homeostasis is established and maintained throughout the cell cycle remains to be discovered. Here, we developed a new technique based on quantification of histone levels to monitor cell cycle progression in individual yeast cells with unprecedented accuracy. Our analysis establishes the existence of a strong mechanism controlling bud size in G2/M that prevents premature entry into mitosis, and contributes significantly to the overall control of size variability during the cell cycle. While most G1/S regulation mutants do not display any strongly impaired size homeostasis, mutants in which B-type cyclin regulation is altered display large cell-to-cell size variability. Our study thus demonstrates that size homeostasis is not controlled by a G1-specific mechanism but is likely to be an emergent property resulting from the integration of several mechanisms, including the control of cyclin B-Cdk activity, that coordinate cell and bud growth with division.

## Introduction

Proper cell function requires an appropriate balance of the relative size of cellular organelles. To ensure cell size homeostasis, cells must coordinate growth and division during the mitotic cycle. During the 1970s, genetic studies aimed at deciphering the biochemical architecture of the cell cycle emerged concomitantly with efforts to characterize cell size control in fission (1, 2) and budding yeast (3, 4). In a free-running cell cycle oscillator model (i.e. in the absence of any coupling to control signals, such as cell size), the cell division time is set by the sum of fixed intervals associated with successive cell cycle events (referred to as “Timer”). In this case, the absence of coordination between the cell cycle engine and cell growth may induce deleterious fluctuations in cell size. In contrast, a “Sizer” mechanism has been shown to operate in yeast: the transition to a given cell cycle phase (resp. mitotic entry in fission yeast and DNA replication in budding yeast) occurs when cells have attained a critical size during the preceding phase (resp. G2 phase in fission yeast and G1 in budding yeast) (1, 2). In this case, small cells experience a size-dependent cell cycle delay and therefore a compensatory mass addition that works as a counteracting force to restore size equilibrium.

In the last 10 years, several important advances have been made in unraveling the molecular mechanism(s) responsible for this “size checkpoint,” which transmits cell size information to the cell cycle control machinery. In fission yeast, it was proposed that the polarity protein kinase Pom1 localizes to the cell tips and indirectly inhibits the cyclin-dependent kinase Cdk1, allowing the cell to sense its elongation and therefore control mitotic entry (5). A later study invalidated this model by showing that Pom1 deletion does not alter size homeostasis, as would be expected following disruption of a core player in the size signaling pathway (6). Further work proposed that Cdr2, a target of Pom1, is responsible for the coupling between cell geometry and cell cycle progression (7). However, this hypothesis has not yet been validated by measuring the size-compensation mechanism in the corresponding mutant background (i.e., *cdr2Δ*).

Early models of cell size regulation in G1 in budding yeast proposed that the commitment (called “Start”) to an irreversible round of division in response to cell growth is controlled by the cyclin Cln3, which is a key regulator of G1 progression and the function of which might be coupled to cell size in various ways (8, 9). Alternatively, recent work showed that the concentration of Whi5, a major inhibitor of G1/S cyclin expression, gradually decays during G1 but is synthesized in a cell size-independent manner during S/G2/M phases; thus, the nuclear concentration of Whi5 is larger in small daughter cells compared with the large mother cells at birth (10). According to this model, coupling between cell growth and cell cycle progression in G1 originates from cell size-dependent dilution of this G1/S inhibitor. Nevertheless, although the *WHI5* mutant displays a small cell size phenotype (11), the G1 size-compensation effect is reduced but not abolished (12, 13), and the overall width of the cell size distribution of Whi5 mutants and wild-type (WT) yeast are similar (11). As with Pom1 in fission yeast(6), the contribution of Whi5 to the overall size homeostasis in budding yeast therefore remains a matter of debate.

*whi5*Δ mutants and cells carrying other genetic perturbations that induce a premature G1/S transition also display compensatory growth in S/G2/M (14-16), which is analogous to the “cryptic” G1/S size control observed long ago in *wee1Δ* fission yeast (17). These observations suggest that, unlike other cell cycle checkpoints (e.g., DNA damage) in which a single sense-and-signal machinery controls cell cycle progression, cell size homeostasis may be maintained by multiple mechanisms that cooperate to coordinate cell growth and division throughout the entire cell cycle. Adding further complexity, previous work has shown that the magnitude of the size-compensation effects during G1 is greatly affected by mutation of several genes with no direct link to G1/S signalling (14). This indicates that size control may result from a complex interplay between the regulatory mechanisms involved in cell cycle progression.

Recent observations in bacteria proposed that a size-compensation mechanism may not even be necessary to ensure cell homeostasis. In contrast to a Sizer mechanism, in which cell size variation during the cell cycle is negatively correlated with the initial cell size, bacteria passively reach size homeostasis through an “Adder” mechanism, whereby a constant amount of cellular material is added at every cell cycle (18-20). However, as recently analyzed in budding yeast, despite the existence of a clear “Sizer” in G1, the effective size control mechanism during the whole cell cycle may be perceived as an “Adder”(13, 18), further raising the question of the integration of multiple size regulation steps during cell cycle progression (21).

By restricting the focus to the G1 size control mechanism, most previous studies overlooked the existence of other size control mechanisms at other cell cycle stages, and, *a fortiori*, how they are integrated to ensure the overall size homeostasis throughout the cell cycle. This is in part because, unlike G1 and mitosis, others phases of the cell cycle could not be accurately resolved in single cell measurements. Therefore, a global quantitative analysis of size compensation effects during the entire cell cycle is required to determine how each phase contributes to cell size control, and how this is perturbed in mutants of cell cycle regulation. Furthermore, the strength of size control was usually assessed by simply measuring the magnitude of size compensation effects, but ignoring how the actual cell size variability - which is the key marker of size homeostasis evolves during the cell cycle. Last, mutants in which the overall size homeostasis - and not only G1 compensatory growth - is truly impaired remain to be identified, which is decisive to improve our understanding of the genetic determinism of size control.

To address these deficits, we have developed a new microscopy technique based on real-time measurements of histone levels to monitor the successive phases of the cell cycle in individual cells in an automated manner. This methodology allowed us to measure a large number of cell cycle phase and cell size-associated variables in 22 mutants, totaling up to 15,000 cell cycles per genotype. Using this dataset, we quantitatively established the existence of a compensatory growth mechanism operating on the bud size during G2 in WT cells, thus confirming the existence of multiple size-dependent inputs in size control (16, 22), in agreement with theoretical predictions (23), and clearly ruling out the “cryptic” nature of size control in G2. Unexpectedly, among the cell cycle genes tested that affect size compensation in either G1 or G2, we found that genes related to the regulation of cyclin B-Cdk activity had the strongest impact on size homeostasis. This finding contrasted with mutants of G1 regulators, which displayed only modest effects on size control. Quantification of cell size variability during the cell cycle showed that phase-specific compensatory growth directly controls the noise strength in cell size distribution, as demonstrated using a linear map model that accommodates experimental data presented in this study. Therefore, unlike the prevailing model of a dominant G1-specific size control checkpoint, our analysis reveals that cell size homeostasis results from the integration of multiple interdependent elements acting at different stages of the cell cycle on different cellular compartments.

## Results

### A New Technique to Monitor Cell Cycle Progression in Live Yeast Cells

To obtain a precise assessment of cell size control during cell cycle progression, we sought a quantitative marker of the successive cell cycle phases in individual growing cells. Studies to date have relied on monitoring of bud emergence or of a fluorescent budneck marker, neither of which can distinguish between S, G2, and M phases. We reasoned that the burst of histone synthesis could serve as an accurate marker of S phase, thanks to the tight reciprocal coupling of DNA replication and histone synthesis, which has been characterized in detail (24-27). Therefore, determining the onset and the end of the burst of histone expression would allow us to extract the duration of S-phase, but also deduce the duration of phases that precede (G1) and succeed (G2/M) DNA synthesis.

To this end, we took a strain carrying a fluorescent protein cassette fused to one of the histone 2B loci (*HTB2*, Fig. 1A), which has been extensively used as a nuclear marker. We used a superfolder GFP (sfGFP) protein to ensure a fast maturation of the fluorophore, in order to prevent artifacts in measurements of histone dynamics, as described below. We monitored yeast cell growth in a microfluidic device which allowed us to track the successive divisions of individual cells forming bi-dimensional microcolonies (See Methods and Fig. 1 and S1), as previously described (28).

**Fig. 1.**
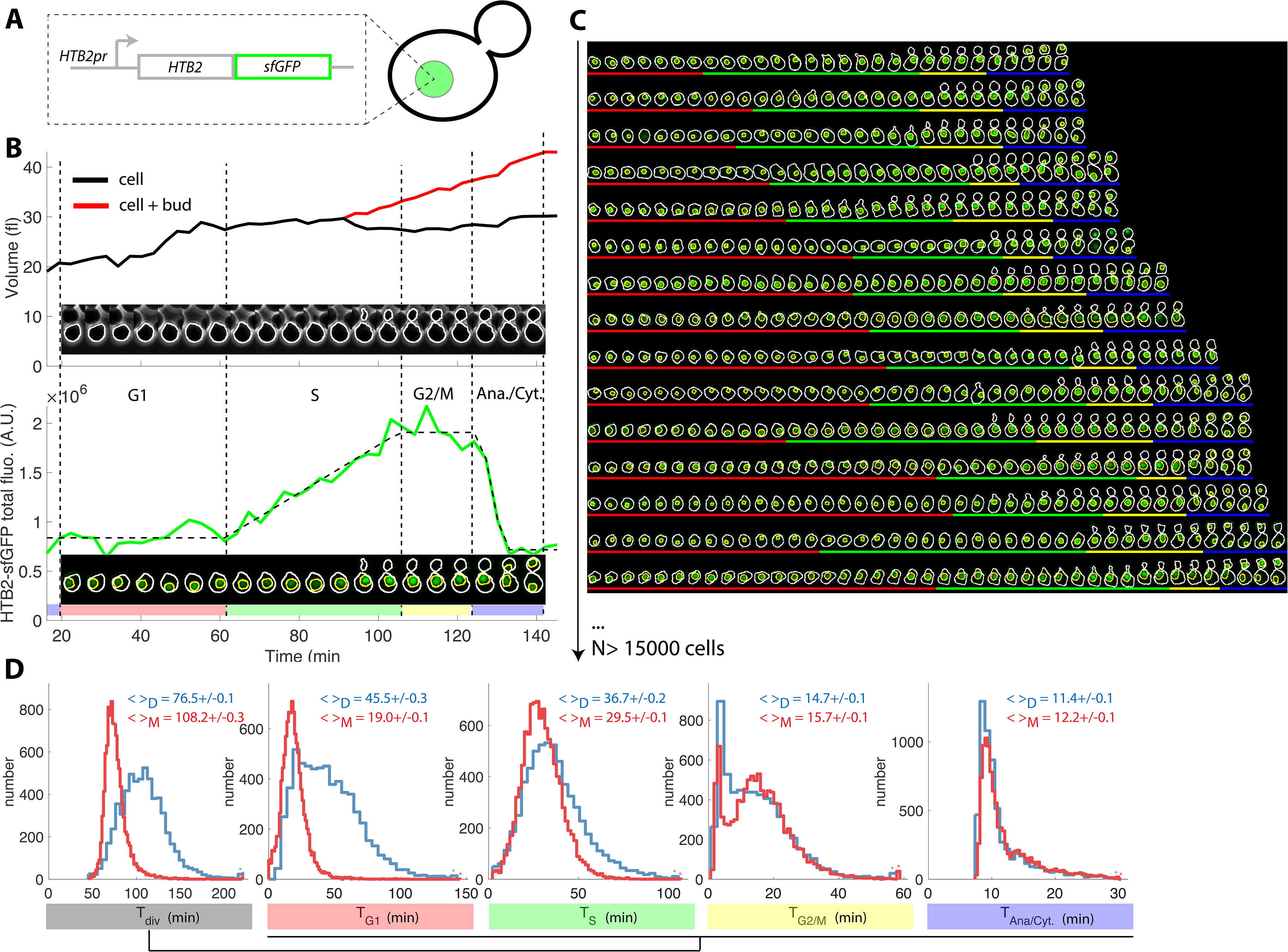
Tracking of cell cycle phases in individual cells. (A) Principle of the H2B-GFP fluorescence marker used track cell cycle progression (B) Sequence of phase contrast (upper) and fluorescence (lower) images of a sample wild-type daughter cell carrying a histone marker (HTB2-sfGFP), displayed with a 6-min interval. Segmented cell and nuclear contours are indicated in white and yellow, respectively. The upper and bottom panels show the quantification of cell (and bud) volume and total fluorescence signal (green curve) over time, respectively. The dashed line shows the best fit of a piecewise linear model to the fluorescence signal, which is used to segment the cell cycle into distinct phases (see text for details), as indicated using a specific color code. Vertical dashed lines highlight cell cycle phase boundaries. (C) Sample dynamics of 15 individual daughter cells during one cell cycle. The green signal represents nuclear fluorescence of the HTB2-sfGFP marker. White and yellow lines indicate cellular and nuclear contours, respectively. Colored segments (G1, red; S, green; G2/M, yellow; anaphase/cytokinesis, blue) indicate cell cycle intervals, as determined using the procedure described in (B); (D) Histogram of durations of cell cycle intervals and overall cell cycle for WT daughter (D; N=6079) and mother (M; N=10775) cells. The legend indicates the mean +/- standard error on mean.

Plotting total nuclear *HTB2*-sfGFP fluorescence as a function of time over one cell cycle (Fig. 1B, see Fig. S2A–D and Supporting Information for details of all quantifications), revealed a fluorescence plateau during the unbudded period of the cell cycle, followed by a linear ramp starting shortly before budding, and a plateau during the budded period of the cell cycle. This pattern was terminated by a sudden drop in fluorescence, corresponding to the onset of anaphase and nuclear division.

Quantification of fluorescence levels gave a consistent ~2-fold enrichment in histones before compared with after the histone synthesis phase (Fig. S2E and F), as expected by the doubling of DNA content. Similarly, we verified that HTB2-sfGFP fluorescence was evenly partitioned between mother and daughter cells upon nuclear division (Daughter/Mother [D/M] asymmetry = 0.94 ± 0.01 Fig. S2G).

To further check that the measurement of total histone content over time is a reliable and physiological way to score cell cycle progression in individual cells, we performed a series of control experiments. First, we compared the division time (by measuring the anaphase to anaphase interval) of a strain carrying a constitutive NLS-GFP marker with a *HTB2*-GFP strain. We observed that the GFP-tag at the *HTB2* locus only modestly affected cell division (Figure S3A-C). Of note, unlike the piecewise expression pattern observed with the *HTB2*-GFP strain (Figure S3A), the NLS-GFP strain yielded a continuous increase in total fluorescence throughout the cell cycle, as expected by a constitutive marker.

Next, we investigated how the maturation time of the fluorescent reporter affects the ability to accurately monitor the burst in histone level during S-phase. For this, we followed the expression of a second histone H2B marker, *HTB2*-mCherry, over time. Importantly, only a linear ramp followed by the anaphase drop could be discerned, in striking difference with the pattern observed with sfGFP (compare Fig. S4A and B with Fig. S4C and D). A numerical model confirmed that this effect could be quantitatively explained by the much longer maturation time of mCherry (~45 min (29)) compared with sfGFP (~5 min (30)), which blurs the apparent dynamics of histone synthesis (Fig. S4E–H).

Although histone levels monitoring provides the timings of S phase and nuclear division, cytokinesis cannot be timed and, therefore, the duration of G1 cannot be deduced. To circumvent this problem, we used the septin subunit Cdc10-mCherry fusion as an additional cytokinesis marker (Fig. S5A–C). We measured that cytokinesis (the sudden drop in Cdc10-mCherry fluorescence) and nuclear division were tightly correlated (Pearson coefficient 0.94), with a median offset of 5.6 ± 0.4 min between both events (Fig. S5D and S5E). Therefore, for the sake of simplicity, we chose to ignore cell-to-cell variability in this part of the cycle and, in the rest of the paper, we arbitrarily defined cell cytokinesis as an event occurring 5.6 min after the end of anaphase.

Similarly, we used the Whi5-mCherry fusion protein to assess the coordination between cell cycle Start (as defined by nuclear exit of the transcriptional repressor Whi5) and the onset of histone synthesis (Fig. S5F–H). Start consistently occurred before the onset of histone synthesis (Fig. S5I–J), which was expected because *HTB2* expression is controlled by the G1/S-specific transcription factors SBF/MBF. Taken together, these results confirmed the tight coordination between cell cycle progression and our measurements of the dynamics of histone expression.

To extend this preliminary analysis, we developed custom MATLAB software *Autotrack* to automate the processes of cell and nucleus contour segmentation, cell tracking, histone content measurement, and mother/daughter parentage determination (Fig. S6 and Supporting Information). We then used a piecewise linear model to identify the histone synthesis plateaus and ramp in the raw data, which allowed us to extract 4 intervals per cell cycle (Figure 1B-C and Movie S1): G1 (plateau), S (linear ramp), G2/M (plateau preceding anaphase), and the interval between anaphase onset and cytokinesis (referred to as “Ana”), taking into account our hypothesis that the period between the end of anaphase and cytokinesis was constant, as mentioned above.

Using this method, we extracted the duration of cell cycle phases for up to ~500 cells in each of the 8 cavities in each independent chamber. By pooling 17 replicate experiments, we collected ~26,900 cell cycles for WT cells (Fig. 1C) of which 63% passed our quality control procedure aimed at discarding cells with segmentation/tracking or data fitting issues (see Supporting Information). To decrease the rate of cell rejection due to noise in histone level signals, we tested multi-z-stack acquisition for HTB2-sfGFP fluorescence. However, this only marginally improved the signal to noise ratio (Fig. S7A–C) while greatly affecting the cell cycle duration likely due to photo-damage (p<0.001, Fig. S7D). Therefore, we retained the single plane acquisition method.

Using this analysis, we found that the cell cycle durations for WT cells were in good agreement with data obtained using other markers or methodologies. Thus, the durations for mothers and daughters, respectively, were: G1 (19.0±0.1 and 45.5±0.3min) (31), S (29.5±0.1 and 36.7±0.2 min) (32), G2/M (15.7±0.1 and 14.7±0.1 min), and Ana (11.4±0.1 and 12.2±0.1 min), Fig. 1D and Table 1. Importantly, the large sample size allowed us to identify statistically significant differences in these intervals. For instance, the S phase was 7.2 min longer in daughters compared with young mother cells (p < 0.001; Fig. 1D). In addition, this interval converged toward an asymptotic value over several divisions following cell birth (Figure S8). This contrasts with G1 duration, which decreased abruptly when daughters (replicative age 0, Fig. S8) became mother cells (replicative age > 0, Fig. S8) in their subsequent division, and G2/M, the duration of which is quite independent of the replicative age of the cells. This phenomenon explains the previously reported (33) *progressive* shortening of the cell cycle duration with the replicative age of the cell (see also Fig. S8) and illustrates the power of our technique to quantitatively measure the temporal distribution of cell cycle intervals.

### Effects of Environmental and Genetic Perturbation on the Distribution of Cell Cycle Phase Durations

Because our methodology allowed us to detect even minor differences in cell cycle duration, we sought to validate its robustness by measuring the timing of cell cycle phases following perturbation by diverse environmental and genetic perturbations approaches that have been extensively studied using other techniques.

First, we asked whether our methodology could capture the lengthening of the S phase induced by hydroxyurea (HU), which inhibits DNA replication. As expected, we observed a progressive HU concentration-dependent increase in S phase duration in both mother and daughter cells, from an average of 27.1±0.4 min and 35.1±0.8 min at 0 mM to 48.0±1.4 and 43.9±1.4 min at 50 mM HU, respectively (p<0.001, Fig. 2A). This prolongation of S phase was accompanied by a parallel doubling in G2/M duration (from ~15min to ~30min in both mothers and daughters, p<0.001, Fig. 2A), which is likely due to the activation of the checkpoint that follows DNA damage(34). Interestingly, mothers, but not daughters, exposed to HU experienced a dose-dependent slowing of the entire cell cycle, due to an apparent compensatory decline in G1 duration in the daughters (Fig. 2A).

**Fig. 2.**
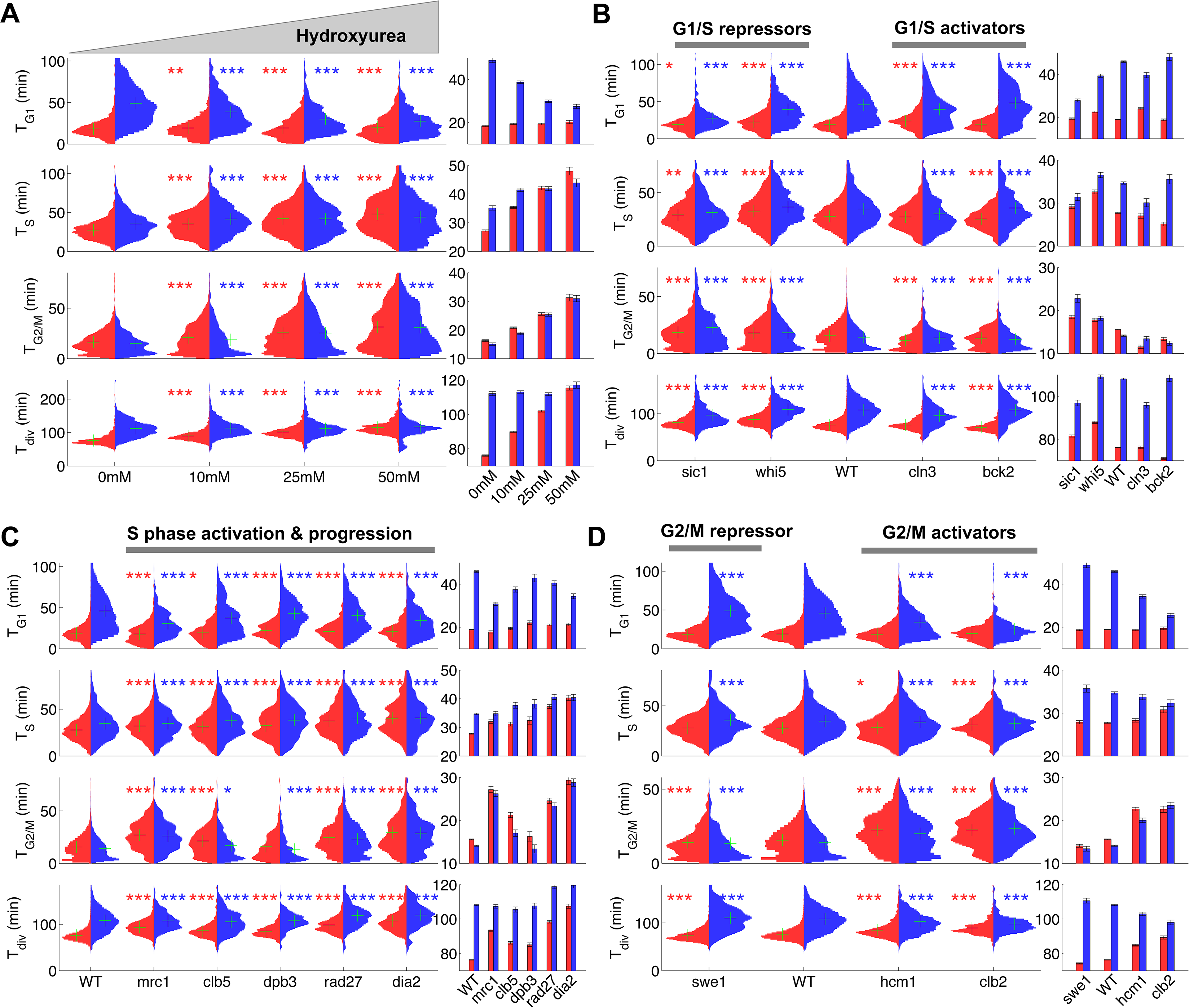
Duration of cell cycle phases in hydroxyurea-treated wild-type cells and in cell cycle mutant strains. (A) Left: Violin plots of the distribution of duration of G1, S, and G2/M phases and total division time (T^div^) for mother (red) and daughter (blue) wild-type (WT) cells at the indicated hydroxyurea concentration. Green crosses indicate the distribution median. Star symbols indicate results of a Kolmogorov-Smirnov test: \**P* < 0.05, \*\**P* < 0.01, \*\*\**P* < 0.001 vs. 0 mM control. Right: Means of the distributions shown on the left. Error bars display standard error on mean (B-D) Same representation as in (A) analyzing various cell cycle mutants (untreated), using the WT strain as a reference for statistical tests.

We next measured the duration of cell cycle phases in cells carrying mutations in important regulators of G1, S, or G2/M phases. First, we confirmed that *sic1* and *whi5* (G1/S transition repressors) daughter cells underwent premature entry into S phase (i.e., shorter G1 duration) compared with WT cells(14), which was accompanied by a compensatory increase in G2/M duration (Fig. 2B)(14). Conversely, mutation of G1/S transition activator *BCK2*(14), but not *CLN3*, caused a small delay in G1 of daughter cells. A slight decrease in G2/M duration was also observed in both *BCK2* and *CLN3* mutants compared with WT cells (Fig. 2B and Table 1).

Next, we monitored the effect of mutations in genes involved in several biochemical pathways related to S-phase (Mrc1, Clb5, Dpb3, Rad27, Dia2). These mutations had previously been shown to induce an abnormally long S-phase interval using a cytometry assay (35). Our results confirmed this observation in all mutants, and, with the exception of *dia2*, the ordering of mutants according to S-phase duration was similar to Koren *et al.* (Fig. 2C)(35). In addition, most of these mutants also displayed a longer G2/M phase, similar to the effect of HU treatment, with the exception of *dpb3* cells (Figure 2A and C). This result suggests that the increased G2/M duration observed following HU treatment and in most S phase mutants is likely to be biological in origin rather than an artefact of our methodology. In this regard, a similar delay in G2/M progression was previously reported in *dia2, mrc1*, and *rad27* mutants, but not in *dpb3* mutants (35).

Lastly, we measured the cell cycle duration in mutants of G2/M progression. We found that deletion of *SWE1*, an inhibitor of mitotic entry, did barely affect G2/M duration, yet induced a slight increase in G1 duration of daughter cells, as previously observed (16). However, mutants defective in Hcm1, a forkhead transcription factor that regulates late S phase genes, or Clb2, one of the main mitotic cyclins, both displayed longer G2/M phases (Fig. 2D), in agreement with previous measurements (36, 37).

Collectively, these results obtained in various mutant backgrounds further establish proof-of-principle for our methodology, in which a single fluorescent marker enables simultaneous measurements of key events associated with cell cycle progression. In total, we monitored the dynamics of cell cycle progression of 22 mutants. The raw cell cycle data are available on a dedicated server (charvin.igbmc.science/yeastcycledynamics/) that allows detailed data exploration and on-the-fly statistical analyses (see Supporting Information).

### Control of Mitotic Entry via a Bud-Specific Size Compensatory Mechanism

Our ability to measure the duration of specific cell cycle phases provides a unique opportunity to investigate in detail the coordination of growth and division during each phase of the cell cycle. We extracted 15 variables (e.g., phase durations, bud/cell volumes and growth rates during unbudded and budded period; see Supporting Information) describing cell cycle progression in both mother and daughter cells. Cell volumes were computed from segmented cell contours assuming an ellipsoid model.

Using this dataset, we sought to identify novel compensatory effects reflecting the existence of size control mechanisms. For this, we systematically measured the Pearson correlation coefficient for all measured distributions of variables in mothers and daughters (Fig. 3A). This analysis successfully confirmed classical results, such as the negative correlation between size at birth (V_birth_) and the duration of G1 (T_G1_) in daughter cells (red stars in Fig. 3A) (31), which is indicative of a G1 compensatory mechanism in small daughter cells, as well as the positive correlation between the growth rate in unbudded cells (μ_unb._) and the cell volume the end of G1 (V_G1_) in daughters cells (blue star in Fig. 3A)(38).

**Fig. 3.**
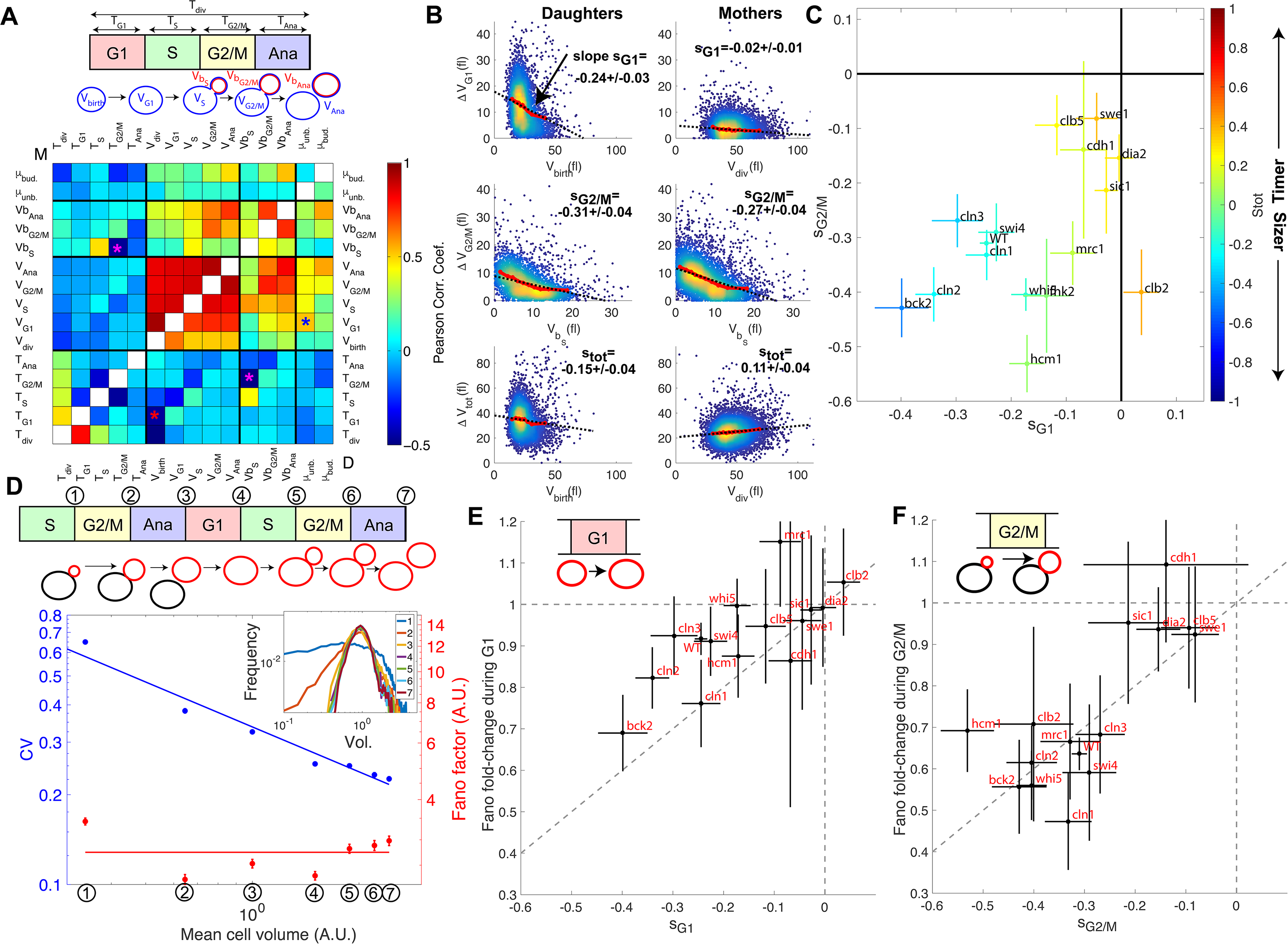
Measurement of size compensation and noise. (A) Top: Schematic of cell cycle phases and definitions of variables used in the correlogram below. Bottom: Each color-coded square in the correlogram represents Pearsons correlation coefficient obtained from the scatter plot associated with the two variables. As indicated on the color scale, blue indicates a negative correlation and therefore highlights the presence of a potential compensatory mechanism. T indicates the duration (min) of each cell cycle phase; V and Vb indicate the mother and bud volumes at each cell cycle phase, respectively. μunb and μbud are the linear growth rate during the unbudded and budded period of the cell cycle, respectively. Ana indicates anaphase to cytokinesis interval. M (top left triangle) and D (bottom right triangle) represent the analyses performed in mother and daughter cells, respectively. Colored asterisks indicate squares of specific interest (see Main text). (B) Scatter plots showing variations in mother/bud volumes at the indicated cell cycle stages. Color indicates point density. Red line shows binning of the scatter plot along the x-axis. Dashed black line is a linear regression through the cloud of points, and the indicated slope (s) represents the strength of the size-compensation mechanism. Error bars (SE) were obtained by bootstrap analysis on the residuals of the linear regression. (C) Strength of size compensation during G1, G2/M, and the entire cell cycle in the indicated mutant backgrounds, calculated as described for (B). The cross color indicates the overall compensation size during the entire cell cycle, as indicated by the color scale. Values of −1 and +1 correspond to an ideal Sizer and Timer, respectively. Errors of the mean were obtained by bootstrap analysis. (D) Measurements of cell and/or bud size variability at the cell cycle phases indicated in the top schematic with a red contour. Left axis represents the coefficient of variation (CV, blue symbols and line) and right axis indicates the Fano factor, as defined in the text (red symbols and line). Inset shows the normalized distribution of cell and/or bud size obtained at different cell cycle stages corresponding to the top schematic. Errors of the mean were obtained by bootstrap analysis. (E) Ratio of the Fano factors at the end and the beginning of the G1 phase for the indicated mutants as a function of the magnitude (slope) of the size-compensation mechanism, as defined in (B). Errors of the mean were calculated using a bootstrap test. The dashed line has a slope 1 and coincides with the point (0;1). (F) Same as in (E), except applied to bud growth during the G2/M phase.

However, this analysis also revealed that the duration of G2/M (T_G2/M_) varies inversely with the volume of the bud at the end of S phase (Vb_S_) in both mother and daughter cells, see magenta stars in Fig. 3A. This indicates that small-budded cells experienced a bud size-dependent delay before entering anaphase. Interestingly, this phenomenon is in agreement with the bud morphogenesis checkpoint model, which proposed that bud growth perturbations lead to a cell cycle arrest that prevents a potentially deleterious entry into mitosis (16). It also matches the conclusion of a recent theoretical model of cell cycle control, in which bud size control appears to play an important role for the overall cell size homeostasis (23). However, another study provided evidence that this control mechanism does not operate as a bud size controller during an unperturbed cell cycle (39). Therefore, to characterize this potential G2 bud size control further, we sought to determine the magnitude of compensatory growth effects during this phase of the cell cycle, and to compare it to the one of other phases. To this end, we monitored variation in cell volume ΔV during G1, G2/M, and the complete cell cycle as a function of initial cell volume (Fig. 3B), according to a methodology widely used in previous studies (18): for an ideal Sizer, the variation in cell volume is such that the final volume V_f_ is constant, independently of the initial one V_i_. In this case, since ΔV= V_f_- V_i_, plotting ΔV versus V_i_ yields a linear relationship with a slope −1. For an ideal Timer (in which the duration of the phase is constant), the slope becomes +1, assuming an exponential growth model(18). Last, an Adder is such that the amount of added volume is independent of the initial volume; in this case, the slope is 0. Therefore, measuring the slopes *s* of ΔV vs. V_i_ plots provide a quantitative assessment of the magnitude of size compensation effects, as well as their deviation from theoretically ideal behaviors (i.e. Sizer, Adder and Timer).

The analysis captured the well-characterized daughter-specific Sizer in G1 (Fig. 3B: slope s_G1_ is negative in daughters, −0.24 ± 0.03, but approaches zero in mothers, 0.02 ± 0.01) (31). It also confirmed that the daughter and the mother cells behave as a weak Sizer and Adder, respectively, over the entire cell cycle (stot = −0.15 ± 0.04 for daughters and 0.02 ± 0.01 for mothers) - in previous studies, the low absolute values of stot showed that the budding yeast cell cycle behaves as an Adder, as observed in other unicellular organisms (13,18).

In addition, the data clearly indicated the existence of a bud-specific size-compensatory growth in G2/M in both daughter and mother cells (s_G2/M_ = −0.31 ± 0.04 and −0.27 ± 0.04 in daughters and mothers, respectively, Fig. 3B). Importantly, we found that the magnitude of the Sizer was strongly reduced when considering the total cell volume (rather than only the bud, see Fig. S9B), or when measuring the magnitude of size compensation during the whole budded period of the cell cycle (Fig. S9C). This very likely explains why G2/M size control has been largely ignored in budding yeast and has been considered as “cryptic”. Instead, our measurements revealed that the G2/M bud size control is of comparable magnitude to the long-known size control in G1. Therefore, these results suggest that several mechanisms may act coordinately to ensure size homeostasis throughout the cell cycle (14).

### Impaired Size Control in Mutants of Cyclin B Regulation and Function

To better characterize the molecular basis of size compensation effects throughout the cell cycle, we measured the magnitude of compensatory growth for daughter cells (using ΔV vs. V plots, as in Fig. 3B) in a subgroup of the cell cycle progression mutants examined earlier (Fig. 2). We found that deletion of the repressor of G1/S cyclin Whi5 decreased the magnitude of G1 control (s_G1_ = −0.17 ± 0.04) but compensated with a slight increase in G2/M control (s_G2/M_ = −0.40 ± 0.06, Fig. 3C). However, these changes were relatively modest, and the overall size-compensation slope was similar in *whi5* and in WT cells (s_tot_ =-0.15± 0.04 and s_tot_=-0.16± 0.09, respectively; Fig. 3C). Deletion of other G1/S regulators (*cln1, cln2, cln3, swi4*) lead to similar conclusions. However, G1 (and, marginally, G2/M) size compensation were slightly improved by deletion of the activator of G1/S transition *BCK2* (s_G1_ = −0.40 ± 0.10, s_G2/M_ = −0.43 ± 0.10), and the overall compensatory growth was stronger than in WT cells (s_tot_ = −0.48 ± 0.13).

In striking contrast to these G1/S regulators, deletion of other cell cycle control genes related to the control of B-type cyclin function, such as *sic1, swe1, clb5*, and *clb2*, induced a much larger decrease in size compensation in both G1 and (with the exception of *clb2*) G2/M phases, as well as the overall compensatory growth (Fig. 3C). For instance, loss of Swe1, which inhibits Cyclin B-Cdk activty and regulates entry into mitosis, leads to a significant Timer behavior (s_tot_ = 0.38 ± 0.06), in which both G1 (s_G1_ = −0.04 ± 0.04) (14) and G2/M (s_G2/M_ = −0.08 ± 0.04) size compensation were largely abolished. Taken together, these data indicate that the compensatory mechanisms ensuring the control of cell size were strongly affected in mutants linked to the regulation cyclin B-Cdk activity but, unexpectedly, only marginally impaired in mutants of the G1/S control network.

### Effective Cell Size Homeostasis during Cell Cycle Progression

To determine how size G1 and G2/M compensation effects actually impact size homeostasis, we sought to directly assess the evolution of the variability in cell size during cell cycle progression. We measured the coefficient of variation (CV) of the distributions of cell/bud volumes at various points in the cell cycle from bud emergence to the next division of the resulting daughter cell (Fig. 3D upper schematic). We found that the CV gradually decreased as a function of cell cycle progression and cell size (Fig. 3D and inset), roughly following a square-root dependency: CV = F^1/2^ / <V>^1/2^, where F is a constant and <V> is the mean cell volume at a given point in the cell cycle (Fig. 3D). This scaling relationship between CV and cell size is to be expected, according to the Central Limit Theorem, assuming that cell growth is the sum of elementary stochastic processes: growth fluctuations tend to average out in larger cell compartments compared to smaller ones. Therefore, to characterize the intrinsic variability in cell size during cell cycle progression and to facilitate comparison among mutants of diverse sizes, we evaluated F (known as the Fano factor (40)), rather than the CV, because F provides a size-independent measurement of noise in cell size during the cell cycle (F is sometimes referred to as noise strength (41)).

As expected, the Fano factor was much more stable than the CV during cell cycle progression of WT cells (Fig. 3D). Still, it displayed some notable variations around the mean at different points in the cell cycle: specifically, F decreased during G2/M and G1 phases, but increased during the rest of the cell cycle, especially during S phase (Fig. 3D). This finding suggested that the Fano factor (i.e cell size noise) may be modulated during cell cycle progression. To check this further, we asked whether the magnitude of the decrease in Fano factor at specific cell cycle phases was consistent with that of the compensatory growth. For this, we plotted the fold-change in Fano during G1 (Fig. 3E) and G2/M (Fig. 3F) for each of the cell cycle mutants. Importantly, we observed that mutants with strong size compensation effects (i.e. with a negative slope) displayed larger reductions in Fano factor in both G1 and G2/M phases. Of interest, the reduction in Fano factor was larger in G2/M (fold-change ~0.65) than in G1 (~0.9) in WT cells (Fig. 3E and F), confirming the importance of cell size control during G2/M. Also, this analysis clearly demonstrates that the magnitude of size compensation mechanisms directly influences size homeostasis in a cell cycle phase-specific manner.

### A Linear Map Model Linking Size Control Efficiency to Size Homeostasis

Following the analysis of compensatory growth during G1 and G2/M, we wondered how the overall (i.e., during a full cell cycle) daughter cell size homeostasis was dependent on the overall size control (s_tot_) in the cell cycle mutants. To explain the quantitative relationship between the magnitude of size control due to compensatory growth and actual variability in cell size, we turned to a noisy linear map of the cell cycle, which provides a simple model to couple phenomenological parameters that describe cell growth and division (42). Under this assumption, the evolution of daughter cell volume V_n_ at the beginning of the cell cycle *n* can be given by (see Supporting Information for detail):

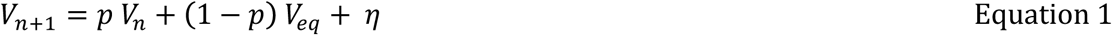

with *p* = *r a*, where *a* characterizes the efficiency of size control (which is directly related to the magnitude of size compensation, represented by the slope *s*_*tot*_ measured in Figure 3B: a = s_tot_+1, see Supporting Information), *r* is the fraction of volume going to the daughter cell at division (asymmetry factor, 0 < r < 1/2), *V*_*eq*_ is the volume of a daughter cell at equilibrium (Fig. 4A), and *η* represents a Langevin noise, such that <*η*> = 0 and <*η*^2^> = constant. Under these assumptions, we demonstrated that the Fano factor would be given by (see Supporting Information for detail):

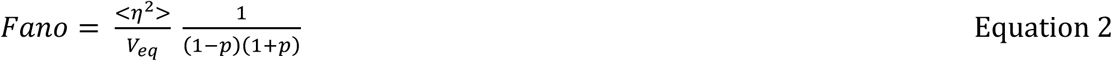

**Fig. 4.**
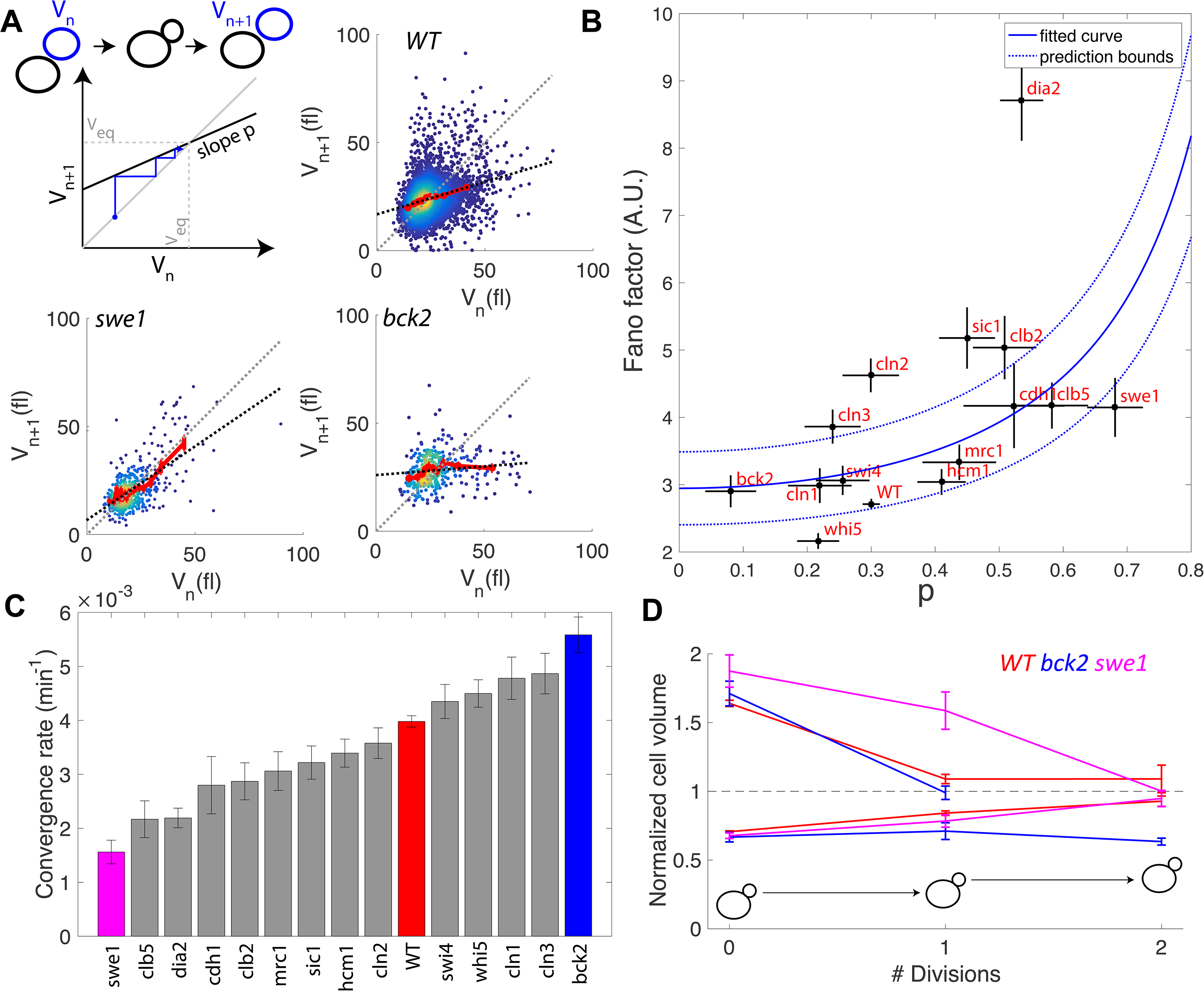
Return map analysis linking cell size noise to the magnitude of the size-compensation mechanism. (A) Top left: Illustration of the return map model, showing the successive iterations of daughter cell size at birth Vn. The size-compensation mechanism can be described by three parameters: steady-state volume Veq, size-compensation strength p, and noise η. The three return maps of experimental data were obtained with wild-type (WT), *swe1*, and *bck2* daughter cells. Color indicates point density. Red line shows binning of the scatter plot along the x-axis. Dashed black line is a linear regression through the cloud of points. The gray dashed line is the diagonal. (B) Average Fano factor during the entire cell cycle as a function of the experimentally measured size-compensation strength p. The black points and bars are the mean ± SEM of the WT cells and indicated mutants. The blue line shows the single parameter fit to the model (see text), yielding the intrinsic noise of the growth process 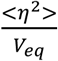= 3.2 ± 0.7, with 99% confidence intervals indicated by the dashed blue lines. (C) Rate of convergence to an equilibrium size for each strain listed in order, based on equation 3 in the main text. Mean ± SEM. (D) Normalized size of successive daughter cells for WT cells and *swe1* and *bck2* mutants, starting from cells that deviate by more than 50% or less than 30% of the equilibrium cell size (indicated by the black dashed line).

This equation indicates that the variability in cell size depends on a, size-independent,intrinsic noise constant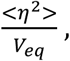 which reflects the stochasticity of the growth process, and an effective size control parameter *p* (with 0 < *p* < 1). Notably, this model predicts a strong non-linearity in size variability as a function of size control parameters, and has two interesting limit cases: for a perfect Sizer (s_tot_ = −1), *p* equals 0, therefore the Fano factor equals the intrinsic noise associated with the growth process, given by <η^2^>/V_eq_,. In contrast, for a perfect Timer (s_tot_ = 1), and assuming symmetrical division of mother and daughters (*r* = *½*), *p* equals 1 and thus there is a divergence in Fano factor, leading to a complete loss of size homeostasis. In the case of budding yeast, which divides asymmetrically (*r*< *½*), such extreme case is impossible. In other words, even with a pure Timer, asymmetrical division is sufficient to limit cell size variability.

To check the validity of this description, we computed the average Fano factor during the cell cycle and calculated *p* by linear regression of single-cell data in WT and mutants (Fig. 4A). We observed large variations in Fano factor among the mutants, which appeared to be correlated with the size control parameter *p* (Pearson correlation coefficient = 0.53, Fig. 4B): overall, Sizers (i.e. with low values of *p*) tend to have less cell size noise than Timers (i.e. high values of p). Of note, this correlation increased if considering *dia2* as an outlier (Pearson correlation coefficient = 0.60). Interestingly, mutants related to the G1/S network (*bck2, swi4, whi5, cln1*, with the exception of *cln2* and *cln3*) generally displayed a noise level comparable to WT and lower than did the mutants associated with the regulation of B-type cyclin function (*clb5, swe1, clb2, cdh1*; Fig. 4B).

Fitting the model prediction to the experimental data (using a single parameter fit <η^2^>/V_eq_) yielded reasonable agreement, despite a large spread in the experimental data and the existence of an outlier, the *dia2* mutant, which failed to fit the model (blue line on Fig. 4B). Therefore, this analysis revealed that the degree of size variability observed in this cohort of cell cycle mutants, associated with diverse roles in cell cycle progression, can be reasonably accounted for by a simple model in which there is a universal noise parameter that characterizes the stochasticity of the growth process, as well as a mutant-specific parameter associated with size control. The deviation of experimental data from the predictions of the model are likely to originate, in part, from the simplistic assumption that size control is a homogenous process throughout the cell cycle, thus ignoring the contributions of size compensation mechanisms in specific phases. In addition, whereas the hypothesis of a linear map was correct for some strains (e.g. WT in Fig. 4A), some deviations were observed in others (i.e. multiple slopes may be needed to describe the behavior of *bck2* in Fig. 4A).

Interestingly, the parameter *p* is related to the rate *λ* of convergence of the linear map, which sets the time it takes for a chain of daughter cells to return to equilibrium following a fluctuation in cell size:

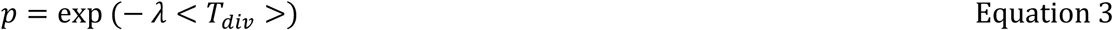

where <T_div_> is the average generation time of a specific mutant. By computing *λ* from p and <T_div_>, we identified a large spread in convergence rates, ranging from 1.5 × 10^−3^ min-1 in *swe1* to 5.5 × 10^−3^ min-1 in *bck2* (Fig. 4C). By selecting the fraction of daughter cells that deviated significantly (larger or smaller) from the equilibrium size at birth and tracking the average size of consecutive daughters, we confirmed that convergence to equilibrium was impaired in the *swe1* mutant but slightly improved in the *bck2* mutant (yet only for large cells, presumably because the slope *p* seems different for small versus large *bck2* cells on Figure 4A, see above) compared with WT cells (Fig. 4D). Therefore, this analysis revealed that mutations of cell cycle regulators not only modify the average duration of specific phases but also control the timescale of size fluctuations and hence the robustness of the cell cycle orbit.

## Discussion

In this paper, we have described a new technique to monitor the duration of successive phases of the cell cycle based on quantification of histone level dynamics in individual growing yeast cells. Most previous single cell analyses tracked cell cycle progression through budding events, ignoring the details of S/G2/M phase events. Our methodology overcomes this limitation and offers new perspectives on the quantification of temporally controlled events in individual cells, such as the coordination between DNA replication and mitosis. Notably, unlike other markers of cell cycle progression (43), our technique is based on a single fluorescent marker, thereby enabling correlative measurements to be made using additional spectrally independent markers.

The large throughput of the image acquisition/processing pipeline developed in our study provides the opportunity to detect mild yet meaningful phase duration phenotypes that were not detected in previous analyses. The discovery that cells replicate their DNA more rapidly with increasing replicative age is a good example of this ability to resolve small differences in cell cycle timing, although this observation needs to be confirmed using complimentary techniques. Since we could not exhaustively analyze the large datasets generated in this study within the scope of this article, we have created a dedicated server to allow further statistical analyses of cell cycle variables in individual cells.

The main interest of our methodology was to enable the identification of size compensation effects throughout the cell cycle in an unbiased approach, and the possibility to assess their role in the establishment of size homeostasis in a quantitative manner. Building on previous studies (16), our work now clearly establishes the link between bud growth and cell cycle progression through G2, and reveals that the magnitude of size compensation is comparable to the well-known G1 size control. Although budding and DNA replication are triggered concomitantly by the activation of the G1/S regulon, the noise in bud size that is observed at the end of S phase suggests that these two processes appear to be largely uncoordinated. Therefore, the function of bud size control during G2/M may be to prevent the potentially deleterious entry of small-budded cells into mitosis following DNA replication. Importantly, this finding challenges the idea of a “cryptic”-type G2 size control in budding yeast, which would only be observed upon appropriate environmental or genetic perturbations. Instead, it supports the hypothesis of universal size control mechanisms across eukaryotes, like fission yeast, in which a G2 size control has long been established (1).

Beyond previous work focusing on the identification of G1-specific size compensation regulators (14), our analysis in mutants broadens our understanding of how the emergence of size homeostasis is connected to the cell cycle control network. Unexpectedly, we found that mutations in activators or repressors of G1 progression had only marginal effects on the overall noise in cell size. In particular, while mutating Whi5 slightly decreased G1 compensatory growth and reinforced G2/M size control, the overall size homeostasis was quite preserved in this mutant. Interestingly, we observed a slight but significant increase in G1 size compensation effects in the *bck2* mutant compared with WT cells, indicating that this phenotype is genetically tunable in both directions.

In contrast, we found that mutations of regulators of cyclin B-Cdk activity had a more pronounced effect on cell size homeostasis: G2/M size compensation was largely abolished in the *swe1* mutant (16), as well as in *sic1* and *cdh1* mutants. Strikingly, all of these mutations also reduced the magnitude of the G1 size control. With a few notable exceptions (*hcm1* or *clb2*), the fact that the magnitude of G1 and G2/M size compensations are somewhat coupled across these mutant strains suggests that enforcing a clear switch in B-type cyclin-Cdk activity between the low (S/G2/early M) and high (late M/G1) APC activity regimes is critical for cell size homeostasis. Further modeling of cell cycle dynamics using a detailed molecular description will be important to clarify this point (44).

A novel feature of our linear map model is the proposed general formula linking the efficiency of cell size control to the noise in cell size, which is reasonably well supported by the experimental data obtained in various mutant backgrounds (Figure 4B). This model predicts a divergence in Fano factor when the behavior of the cell approaches that of an a ideal Timer (19). We speculate that the deletion of some cell cycle genes may render cells inviable, not to loss of an essential biochemical function, but rather to complete loss of size homeostasis, thus impairing the robustness of the cell cycle oscillation. Conversely, even a cell cycle mutant in which stot = +1 (i.e., a perfect Timer according to the previous definition (18, 42)), should be able to control its size if dividing asymmetrically (since p = r (s_tot_ + 1) < 1 when r < ½). Therefore, asymmetric division can be regarded as an additional stabilizer of cell cycle that limits cell size variability.

In conclusion, our study, in which cell cycle progression was monitored with unprecedented accuracy in yeast, demonstrates that size homeostasis does not originate from a G1-specific mechanism, but is likely to be an emergent property resulting from the integration of several mechanisms that coordinate cell growth with division. Our analysis specifically highlights the role of bud size control in limiting cell-to-cell variability, which is likely to be connected to the role played by B-type cyclins in size homeostasis, as identified here. Additional studies connecting further experimental datasets to computational analyses (45) will be instrumental in deciphering how individual components are integrated to ensure size homeostasis throughout the cell cycle.

## Material and Methods

### Strain Construction

All strains were congenic to S288C unless specified otherwise and were constructed following standard genetic techniques. A detailed list of strains is provided in the Supporting Information. HTB2-sfGFP fusion protein was generated by classical PCR-mediated genome editing. Fast maturation of the fluorophore was necessary to ensure accurate determination of the successive cell cycle phases (see Supporting Information). Mutant strains were obtained from the deletion collection of non-essential genes. In the list of constructed strains, we noticed that the *cdh1Δ HTB2-sfGFP* strains was quite unstable and yielded a large fraction of dead cells as well as large multinucleated cells that retained a fast division time and eventually outgrew the rest of the population.

Indeed, the *cdh1Δ* mutation has previously been described to induce genomic instabilities (e.g. chromosome loss, etc..)(46). We hypothesize that introducing the histone marker in this background exacerbates this phenotype. To circumvent this issue, we used freshly thawed cells from frozen stock.

### Microfabrication and Microfluidics Setup

Microfluidic chips were designed and made using standard techniques as previously described (28). The microfluidic devices, which feature 8 independent channels, each consisting of 8 chambers, allow parallel monitoring of 8 genetic backgrounds in the same time-lapse assay. The microfluidic master was made using a standard SU-8 lithography process at the ST-NANO facility of the IPCMS (Strasbourg, France). CAD files and detailed dimensions of the chip are available upon request. The micro-channels were cast by curing PDMS (Sylgard 184,10:1 mixing ratio) and then covalently bound to a 24 × 50 mm coverslip using plasma surface activation (Diener, Germany). Chips were then baked for 1 h at 70°C to improve the sealing between PDMS and glass. Microfluidic chips were connected using Tygon tubing and media flows were driven by a peristaltic pump (Ismatec, Switzerland) with a 30 μL/min flow rate.

### Live Imaging of Yeast

Strains were cultured overnight in synthetic complete medium with glucose and all amino acids. The next morning, the cultures were diluted and allowed to grow until optical density at 660nm reached 0.2−0.5. Each strain was then loaded into independent chambers chosen at random to avoid potential systematic bias in measurements. For each strain, at least two fields of view were recorded during the time-lapse acquisition interval. Each assay included a WT strain as a control. Mutants were analyzed in at least three independent assays. In total, we collected at least 2000 cell cycles per mutant ~25,000 cell cycles for the WT strain.

Cells were imaged every 3 min using an automated inverted microscope (Nikon TI, Nikon, Japan) with a 60× phase contrast objective and a sCMOS camera (Hamamatsu Orca Flash 4.0, Japan) driven by Nikon Software (NIS). Constant focus was maintained using a Perfect Focus system. Fluorophore excitation was performed using LED light(Lumencor X1) and appropriate filter sets. The cells were allowed to grow for up to 10 h in the device, yielding about 500 cells per field of view by the end of the assay.

### Image Processing

Raw proprietary Nikon files (.nd2 format) were converted using Bio-format and bftools packages into MATLAB-compatible lossless jpeg files. We developed custom software (Autotrack; see Fig. S2 and Supplemental Information for details) to: (1) segment and track individual cells in yeast microcolonies, (2) quantify HTB2-sfGFP (or mCherry) levels in individual cells and extract individual cell cycles, (3) determine the duration of individual cell cycle phases, and (4) discard outliers based on specific criteria (detailed in Supplemental Information). Cell volumes were calculated from segmented cell contours assuming an ellipsoid model.

### Dataset Management and Online Data Publishing

All variables extracted during image processing were stored in a mutant-specific database designed to allow straightforward analysis using custom MATLAB software, as described in the Supporting Information. We developed a web application, Yeast Cycle Dynamics, that allows custom statistical analysis of extracted cell cycle data for all mutants in this study. In addition, raw data showing histone levels and cell size as a function of time for all cells can be monitored(charvin.igbmc.science/yeastcycledynamics/). See Supplemental Information for details.

## Acknowledgements

We are very grateful to Damien Coudreuse and Etienne Schwob for insightful discussions and comments on the manuscript, as well as Sophie Quintin and the Charvin lab for careful reading of the manuscript. We thank Sandrine Morlot for help with microscopy. We thank Michaël Knop for sharing plasmids and Joseph Schacherer for the gift of the cell cycle mutants. This work was supported by the ATIP-Avenir program (G.C.), a grant from the Fondation pour la Recherche Médicale (G.C.), and by grant ANR-10-LABX-0030-INRT, a French State fund managed by the Agence Nationale de la Recherche under the frame program Investissements dAvenir ANR-10-IDEX-0002-02.

## Supplementary Figure Legends

**Fig. S1. Principle of the microfluidic device and time lapse experiment**

(A) Photograph and schematics of the microfluidic device used in this study, showing the dimensions of the cavities designed to trap cell microcolonies. (B)Sequence of phase contrast and fluorescence image showing the growth of isolated microcolonies within the microfluidic device. White and yellow contour represent the cellular and nuclear contours, respectively. Cell and nuclear segmentation are described in detail in the supplements.

**Fig. S2. Quantification of H2B-sfGFP fluorescence signal in individual cells**

(A) Typical fluorescence image of a microcolony of cells expressing HTB2-sfGFP. Nuclear contours were generated using a custom segmentation procedure. Scale bar, 5 μm. (B) Higher magnification of the nucleus highlighted in red in (A). Contours of increasing size are used to determine the total HTB2-sfGFP fluorescence and to discard the background signal. (C) Procedure used to remove fluorescence background. Total fluorescence as a function of the nuclear contour size (in pixels). The dashed line is a linear fit to the last 6 points, which represents an asymptotic limit corresponding to background pixels. (D) Total HTB2-specific fluorescence signal obtained after background subtraction. (E) Typical evolution of HTB2-sfGFP fluorescence signal (green line) during a complete cell division cycle. The black dashed line represents a piecewise linear fit to the experimental data. The gray shaded areas represent the interval of G1, G2/M, and subsequent G1 (G1′) phases of the cycle, as defined by the plateaus obtained during the fitting procedure. (F) Distribution of the fluorescence level ratio during G2/M and G1 or G1′. The median ratio is 2.25 ± 0.01. (G) Distribution of the fluorescence level ratio between mother and daughter cell upon nuclear division. The median ratio is 0.94 ± 0.01 (see Methods and Supplemental Information for further details).

**Fig. S3. Influence of the HTB2-sfGFP marker on cell cycle duration**

(A) Top: sequence of phase contrast and fluorescence of cells carrying an NLS-GFP marker at indicated time points. White and yellow line indicate cellular and nuclear contours, respectively. White segment indicates bud/mother parentage. Bottom: quantification of normalized total nuclear Htb2-sfGFP fluorescence during one cell cycle. The dashed lines correspond to automatic detection of anaphase events measured as a sudden drop in total nuclear signal in the mother cell nuclei. Tdiv is the cell cycle duration, as measured from anaphase to anaphase. (B) Same as (A), but with a HTB2-GFP strain. (C) Violin plots showing the distribution of cell cycle duration for mother (red) and daughter (blue) cells with an NLS-GFP or HTB2-GFP marker. Red crosses indicate the distribution median. Star symbols indicate results of a Kolmogorov-Smirnov test: \**P* < 0.05, \*\**P* < 0.01, \*\*\**P* < 0.001, using the HTB2-GFP strain as a reference. Right: Means of the distributions shown on the left. Error bars display standard error on mean.

**Fig. S4. Effect of fluorophore maturation on the apparent dynamics of histone synthesis**

(A) Typical sequence of phase and fluorescence images of a strain carrying a HTB2-mCherry marker. The time stamps are relative to the beginning of the assay. (B) Quantification of the HTB2-mCherry fluorescence signal measured in (A) over multiple cell cycles. The dashed lines correspond to automatic detection of anaphase events measured as a sudden drop in total nuclear signal in the mother cell nuclei. (C and D) Same as in (A and B), but with a strain expressing HTB2-sfGFP. (E-H) Numerical simulation recapitulating the effect of the time delay induced by fluorophore maturation on the measured fluorescence signal following pulsatile synthesis of histones. (E) Schematic of the reaction scheme used to model the expression of a histone-fluorescent protein fusion protein (Htb2-FP, mature; Htb2-FP*, immature). The maturation half-time varies from 5 min for sfGFP to 45 min for mCherry. (F) Pulsatile synthesis rate used to mimic histone synthesis during the cell cycle. (G) Simulation of immature (black line) and mature (red) fluorescent fusion protein according to the model described in (E) following pulses of protein expression as in (F) and assuming periodic cell division events (indicated by dashed lines). In this case, k_maturation_ = ln(2)/45 min-1 (corresponding to the mCherry maturation rate). (H) Same as (G), but with k_maturation_ = ln(2)/5 min-1 (corresponding to the sfGFP maturation rate).

**Fig. S5. Comparison of HTB2-sfGFP fluorescence dynamics with other known cell cycle markers**

(A) Phase contrast and fluorescence images of cells expressing both HTB2-sfGFP and Cdc10-mCherry markers. The white and yellow lines represent the cellular and nuclear contours, respectively. The blue line indicates the contour of the budneck. (B and C) Quantification of total nuclear fluorescence (B) and mean budneck fluorescence (C) as a function of time over multiple cell cycles. The dashed lines indicate the end of anaphase (B) or the end of cytokinesis (C), which are computationally detected as a sudden drop in the fluorescence signal. (D) Histogram of the distribution of the duration between the end of anaphase and the end of cytokinesis, based on automated detection of a sudden drop in the signal (see (C)). (E) Scatter plot showing the time from the end of anaphase to the end of the next cytokinesis as a function of the time from the end of anaphase to the end of the next anaphase (black points). The black dashed line represents the diagonal. The red line is a fit to a line of slope 1, which represents the systematic delay between anaphase and cytokinesis. (F-J) Same analysis as in (A-E), but using Whi5-mCherry as a complementary marker. In this case, both the end of anaphase and entry into S phase are detected (G), as well as the Whi5 nuclear exit (H). In (I), the histogram displays the distribution of time between Whi5 nuclear exit and the onset of histone synthesis. In (J), the time from the end of anaphase to the next phase of histone synthesis is plotted against the time from the end of anaphase to the next Whi5 exit (see Methods and Supplemental Information for further details).

**Fig. S6. Cell segmentation and tracking pipeline**

Procedure for extraction of the duration of cell cycle phases in individual cells. See Supplemental Information for details.

**Fig. S7. Effect of multi-plane acquisition on cell cycle phase quantification**

(A) Sequence of phase contrast and fluorescence images of cells carrying an HTB2-sfGFP marker, based on a 6-image z-stack acquisition (spacing 300 μm). The sequence in the middle and bottom represent the max projection and the image in the middle of the stack, respectively. (B) Quantification of total HTB2-sfGFP fluorescence obtained using max projection (red line) or the image in the middle of the stack (blue line). The dashed line indicates the end of anaphase, detected computationally. (C) Fourier spectrum of the datasets displayed in (B), averaged over more than 400 cell cycles. (D) Left: Violin plots showing the distribution of duration of the indicated cell cycle phase or the entire cell cycle (Tdiv) in a wild-type strain expressing HTB2-sfGFP, acquired using a single plane or a 6-image z-stack. The red and blue plots correspond to mother and daughter cells, respectively. \**P* < 0.05, \*\**P* < 0.01, \*\*\**P* < 0.001 vs. control (single plane) using a Kolmogorov-Smirnov test. The green cross indicates the median of the distribution. Right: Mean of the distributions represented on the left.

**Fig. S8. Evolution of cell cycle duration as a function of replicative age**

Left: Violin plots showing the distribution of duration of the indicated cell cycle cell cycle phase as a function of the replicative age of the cells (Age=0 and Age>0 correspond to daughter and mother cells, respectively). \**P* < 0.05, \*\**P* < 0.01, \*\*\**P* < 0.001 vs. replicative age of 1 using a Kolmogorov-Smirnov test. The green cross indicates the median of the distributions. Right: Mean of the distributions represented on the left. Error bars indicate standard error on mean.

**Fig. S9. Comparison of size compensation effects in the budded part of the cell cycle**

(A) Left: schematic showing the definition of variables used to assess the magnitude of size compensation effects during the cell cycle. Right: scatter plots showing variations in cell/bud volumes during G2/M, as a function of bud volume by the end of S phase. Color indicates point density. Red line shows binning of the scatter plot along the x-axis. Dashed black line is a linear regression through the cloud of points, and the indicated slope (s) represents the strength of the size-compensation mechanism. Error bars (SE) were obtained by bootstrap analysis. Right: same as the scatter plot in the middle, but with G2/M duration. (B) Same as (A), but plotting total variations in cell/bud volumes during G2/M as a function of total cell volume (i.e. cell and bud) at the end of S phase. (C)Same as (A), but plotting total variations in cell/bud volumes during S/G2/M/Anaphase as a function of total cell volume (i.e. cell and bud) at the end of S phase.

**Fig. S10. Evolution of cell cycle duration as a function of replicative age**

Left: Violin plots showing the distribution of duration of the indicated cell cycle cell cycle phase as a function of the replicative age of the cells (Age=0 and Age>0 correspond to daughter and mother cells, respectively). \**P* < 0.05, \*\**P* < 0.01, \*\*\**P* < 0.001 vs. replicative age of 1 using a Kolmogorov-Smirnov test. The green cross indicates the median of the distributions. Right: Mean of the distributions represented on the left. Error bars indicate standard error on mean.

## Supplementary Movies Legends

Movie S1. Cell cycle progression of wild-type cells. Movie showing the overlay of phase contrast and HTB2-sfGFP fluorescence of 16 individual cells during one cell cycle with 3-inute interval. Neighboring cells have been masked for clarity, and images of individual cells were rotated so that the budding axis is identical for all cells. The red line highlights cell and bud contours. The white progress bar on the right indicates the total HTB2-sfGFP signal within the nucleus, which is used as a proxy for progression. The progress bar on the left shows the sequence of cell cycle phases as indicated on the legend.

